# Synchrony in the periphery: inter-subject correlation of physiological responses during live music concerts

**DOI:** 10.1101/2020.09.01.271650

**Authors:** Anna Czepiel, Lauren K. Fink, Lea T. Fink, Melanie Wald-Fuhrmann, Martin Tröndle, Julia Merrill

## Abstract

A concert is a common event at which people gather to share a musical experience. While techniques are increasingly offering insights into naturalistic stimuli perception, this study extended methods to a more ecological context in order to explore real-world music listening within a concert setting. Cardiorespiratory, skin conductance, and facial muscle responses were measured from participants attending one of three concerts with live chamber music performances of works of varying Western Classical styles (Viennese Classical, Contemporary, and Romantic). Collective physiological synchronisation of audience members was operationalised via inter-subject correlation (ISC). By assessing which musical features (obtained via Music Information Retrieval and music-theoretical analyses) evoked moments of high synchrony, logistic regressions revealed that tempo consistently predicted physiological synchrony across all concerts in Classical and Romantic styles, but not the Contemporary style. Highly synchronised responses across all three concert audiences seemed to occur during structural transitional passages, boundaries, and at phrase repetitions. The results support the idea that group synchronisation is linked to musical arousal, structural coherence, and familiarity. By employing physiological ISC and an inter-disciplinary musical analysis, the current study demonstrates a novel approach to gain valuable insight into experiences of naturalistic stimuli in an ecological context.

## Introduction

A concert is a common event at which people gather to share a musical experience, particularly in Western society. Although brain imaging techniques can implicitly measure naturalistic musical perception without behavioural ratings^1–5^, these methods lack applicability in more typical listening situations, particularly those where listening happens in a group, such as in a concert. Not only does a concert provide a naturalistic setting, but live performances can be more immersive^6^, evoke stronger emotional responses^7–9^, while co-presence of audience members can increase physiological and emotional experience^10^. Portable methods such as motion capture^11^ or mobile measurement of the autonomic nervous system (ANS, e.g., cardiology)^10,12^ seem promising in such a setting. As our interest lay in the experience of a Western art music concert, in which listeners are typically still^13^, we focused on ANS measurements during listening of full-length musical works in a naturalistic concert context.

Responses of the sympathetic division of the ANS such as an increased (phasic) skin conductance response (SCR, i.e., sweat secretion), an increase of heart rate (HR), and respiration rate (RR) as well as responses of zygomaticus major (smiling) and the corrugator supercilii (frowning) muscles (electromyography [EMG] activity) typically reflect stress, attention^14,15^, or affective processing^16–19^. In terms of auditory responses, changes in SCR, HR, RR, and EMG activity – reflecting a startle^20–22^ or orienting response^23^ – have been associated with pitch changes^24,25^ and tone loudness (the louder the sound, the greater the SCR amplitude^26,27^) as well as deviations in timbre, rhythm, and tempo^27^. Additionally, physiological responses to music may indicate felt arousal and valence of acoustic features^19,28,29^. For example, faster and increasing tempi are associated with greater arousal^30–34^, increased SCR^27,31,35,36^ and HR^31,35,37,38^, whereas slow-paced (low arousal) music reduces HR and breathing^39,40^. Loudness is positively correlated with arousal^32,41,42^, and, correspondingly, changes in SCR^43,44^ and HR^32,45^. Harmonic ambiguity may also be perceived as arousing^12^, with unexpected chords^46,47^ and notes^12^ (i.e., lower clarity of key) evoking SCR increases. Timbral features, such as brighter tones and higher spectral centroid, are associated with higher arousal^32,48,49^, which correlates somewhat to SCR^35,50^. Importantly, these physiological responses are modulated by musical style: previous studies found that HR increases with faster tempo in Classical music, but decreases with faster tempo in rock music^38^, whereas HR is lower in atonal, compared to tonal music even with both styles controlled for emotion^51^.

Although this research generally supports the idea that physiology is associated with musical features and style, there are many inconsistencies within findings such studies (see^52,53^). This could partly be due to most studies having (too) carefully chosen or constructed stimuli to have little variability in acoustic features (e.g., they use a constant tempo and normalise loudness). While there is value in tightly controlling individual musical features, more research into naturalistic stimuli – which typically has a rich dynamic variation of interdependent features – is required to extend ecological generalisability^54^.

Although previous work on naturalistic music has correlated neural and physiological responses to dynamically changing acoustic features^1–3,50^, or extracted epochs based on information content in the music^12^, perhaps a more robust way to identify systematic responses to naturalistic stimuli is through analysing synchrony of responses^10,55–57^, in particular via inter-subject correlation (ISC, see review^58^). This method – in which (neural) responses are correlated across participants exposed to naturalistic stimuli^59^ – is based on the assumption that signals not related to processing stimuli would not be correlated. ISC research has demonstrated that highly similar responses occur across subjects when exposed to naturalistic films^59–62^, spoken dialogue^63–65^ and text^66,67^, dance^68^, and music^1,5,69,70^, strongly suggesting that highly reliable and time-locked responses can be evoked by (seemingly uncontrolled) complex stimuli (for a review see^71^). Although ISC in fMRI studies can identify region of interests (ROIs) for further analysis (e.g.^59^), ISC can also assess which kind of feature(s) within dynamically evolving stimuli evoke highly correlated responses.

In response to auditory stimuli, higher synchronisation (operationalized via ISC) of participants’ responses is associated with structural coherence, familiarity, and emotional context of stimuli. For example, ISC is higher when listening to original, compared to phase-scrambled, versions of music^69,70^ and spoken text^67^. ISC may also reflect familiarity and engagement: ISC is higher when listening to familiar music, compared to unfamiliar music; however, upon repeated presentation, ISC drops with repetitions of familiar, but not unfamiliar music^5^. Moments of collective synchrony additionally seem to be linked to emotional arousal, where higher correlation coefficients of fMRI^59^, EEG^61^, SCR and respiratation^72^ coincided with moments of high arousal in films, such as a close-up of a revolver^61^, gun-shots, or explosions^59^ as well as close-ups of faces and emotional shakiness in voice^72^. However, our understanding of music and ISC is still in its infancy, and it is unclear which musical features – and at which level or time frame – can evoke synchronised physiological responses, particularly in more naturalistic listening situations outside the laboratory.

In addressing these questions, the current study is – the best of our knowledge – the first of its kind to assess which musical features evoke shared physiological responses during full-length, naturalistic music stimuli in a typical group-listening concert context. To our knowledge, this is additionally the first study to compare how different groups react to the same musical stimuli in such a setting. In light of the replicability crisis^73^, we conducted three identical concerts with different participant groups to test the replicability of the induced physiological responses to music, and thereby, the stability of using a concert hall as an experimental setting. We invited participants to attend one of three instrumental chamber music concerts in an ‘ArtLab’ performance hall (purpose-built for empirical investigations). String quintets by Beethoven (1770-1827), Dean (1961-), and Brahms (1833-1897) with four movements each were performed, showcasing different musical styles (Viennese Classical, Contemporary, and Romantic, respectively) with varying tempo, tonality, compositional structure, and timbre. Common acoustic features associated with physiological responses (as described above) were extracted offline: instantaneous tempo, key clarity, RMS energy (related to loudness^42^), and spectral centroid (timbral feature related to brightness^74–76^), and compared across the styles. Derivatives of these features, representing their degree of change over time, were also obtained. Continuous physiological responses were measured throughout the concert, from which SCR, HR, RR, and EMG activity was extracted from 98 participants (Concert 1 [C1]: 35, C2: 41, C3: 21). For each audience, ISC was calculated over a sliding window (5 musical bars long, i.e., on average 10 seconds long) for each physiological measure, representing the degree of collective synchrony of physiological responses over the time-course of each musical stimulus (see Figure 2a). We identified moments of high synchrony (HS) and low synchrony (LS) from the audiences via upper 20^th^ percentile and 20^th^ percentile centred around *r* = 0. Moments of HS and LS were analysed with respect to the corresponding physiological responses and musical features.

As highly synchronised responses have previously been associated with arousal^59,61,72^, familiarity^5^, or structural coherence^67,70^ in stimuli, we hypothesised that highly correlated physiological responses would be driven by typically arousing acoustic features (higher RMS energy and spectral centroid, faster tempo, and lower key clarity) as well as by compositional structure and familiarity of the different styles. We additionally hypothesized that robust physiological responses to music would be consistent across repeated concert performances.

## Results

### Comparison of performances

As the music was performed by professional musicians, we expected all concert performances to be acoustically similar. We found no significant differences between performances for loudness, tempo, timbre, and duration (see Supplementary Table S1). Correlations of instantaneous tempo, timbre, and loudness between all performances reached *r* > .6, *p* < .001, confirming that they were comparable enough to allow for further statistical comparisons of listeners’ physiology between audiences.

### Stimuli analysis

From exploring the extracted loudness, timbre, tempo, and key clarity, Figure 1a supports the idea that the stimuli offered a rich variation of acoustic features. In comparing styles, contrasts revealed that features of the Contemporary work (Dean) differed acoustically from the Classical (Beethoven) and Romantic (Brahms) styles: Dean had significantly lower RMS energy compared to Brahms (Brahms – Dean for C1: β = 0.005, SE = .002, *p* = .039; C2: β = 0.005, SE = .002, *p* = .037; C3: β = 0.005, SE = 0.002, *p* = .043), significantly lower key clarity compared to Beethoven (Beethoven – Dean: β = 0.08, SE = .02, *p* = .010) and Brahms (Brahms – Dean: β = 0.11, SE = .02, *p* = .001), and significantly higher spectral centroid compared to Beethoven (Beethoven – Dean C1: β = −320.36, SE = 78.70, *p* = .007; C2: β = −277.00, SE = 89.00, *p* = .030; C3: β = −252.46, SE = 77.7, *p* = .024) and Brahms (Brahms – Dean C1: β = −320.90, SE = 79.2, *p* = .007; C2: β = −264.90, SE = 89.4, *p* = .037; C3: β = −251.63, SE = 78.10, *p* = .02). Although tempo did not significantly differ between the styles (all *p* > .375), a noticeable division of tempi distribution in Beethoven and Brahms (see Figure 1a) shows a typical composition practice of contrasting faster and slower movements in Classical/Romantic style.

**Figure 1.**
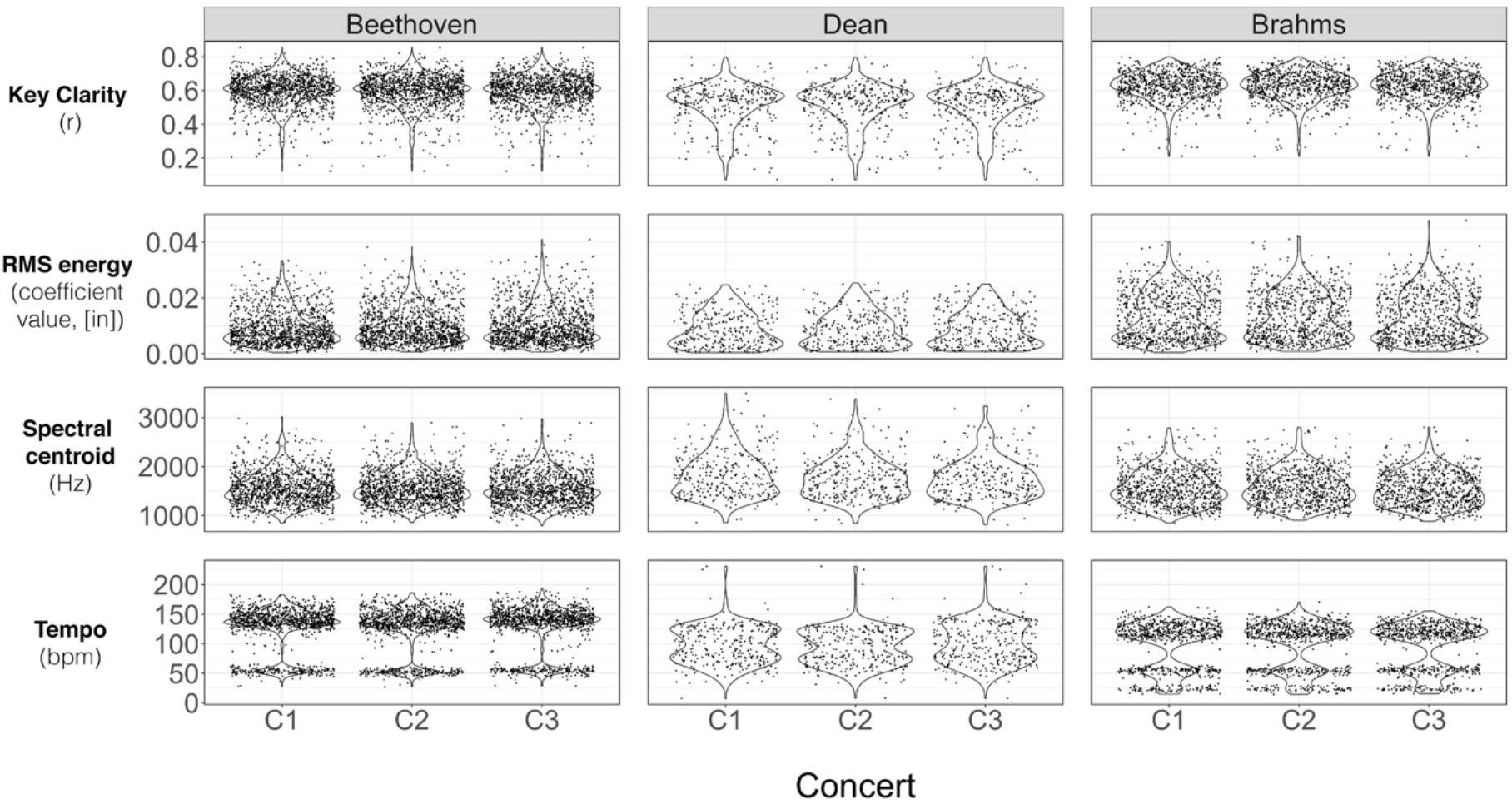
Acoustic features per bar, per piece, per concert. Top to bottom panels show Key clarity, RMS energy, Spectral centroid, and tempo values per bar. Left panels show values for Ludwig van Beethoven (String Quintet in C minor, op. 104, 1817), middle panels for Brett Dean (Epitaphs, 2010), and right panels for Johannes Brahms (String Quintet in G major, op. 111, 1890). Separate violin plots show different concerts.

### Physiological responses at high/low synchrony

As shown in Figure 2b, HR was overall lower at HS bars compared to LS bars (HS – LS C1: β = −0.04, SE = .01; C2: β = −0.06, SE = .01; C3: β = −0.03, SE = .01; all *p* <.002), but HR increased during HS from the onset bar (bar0) to the last bar (bar4) in the correlation window for C1 (bar0 – bar4 in HS: β = −0.05, SE = 0.02, *p* = .031) and C3 (bar0 – bar4 in HS: β = −0.06, SE = 0.02, *p* = .04) (see Supplementary Table S2 for main effects). RR was also significantly lower overall at HS compared to LS bars for C3 (HS – LS: β = −0.03, SE = .008, *p* < .001), but significantly increased across correlation window for all concerts (bar0 – bar4 in HS for C1: β = −0.06, SE = .01, *p* = .003; C2: β = −0.07, SE = 0.01, *p* < .001; C3: β = −0.09, SE = 0.02, *p* < .001). SCRs were significantly higher at HS compared to LS (HS – LS for C1: β = .07, SE = .01; C2: β = 0.03, SE = .01; C3: β = 0.09, SE = .010; all *p* <.001), with a decrease across correlation window (bar0 – bar4 in HS for C1: β = .17, SE = .02; C2: β = .15, SE = .02; C3: β = .18, SE = .02; all *p* < .001). EMG activity was also significantly higher at HS time points in C1 (β = 0.01, SE = .01, *p* = .012) and C3 (β = 0.02, SE = .01, *p* = .012) and decreased across the correlation window in C2 and C3 (bar0 – bar4 in HS for C2: β = .05, SE = .01, *p* > .001; C3: β = .06, SE = .02, *p* > .007). In short, compared to the LS moments, there was higher arousal in HS moments, indicated by higher SCR magnitude and EMG activity, and a general RR and HR increase.

**Figure 2.**
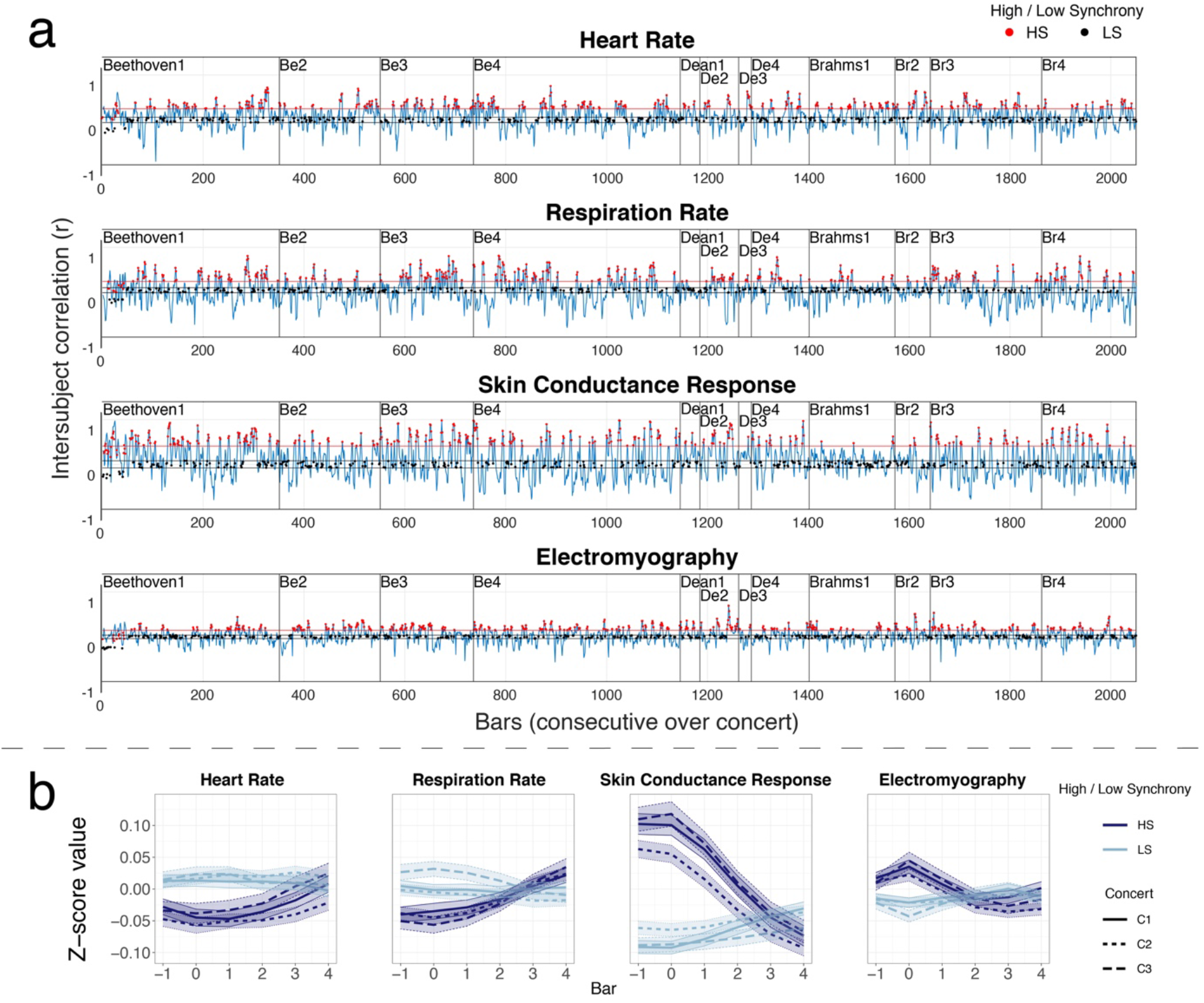
Inter-subject correlation (ISC) across concerts and bars of high- and low-synchrony. **a.** ISC time courses for heart rate (HR, row 1), respiration rate (RR, row 2), skin conductance response (SCR, row 3), and electromyography activity of zygomaticus major muscle (EMG, row 4) for concert 1. Moments of high and low synchrony are marked with red and black dots, respectively. Red lines signify the 20^th^ percentile threshold, while black lines signify the 20^th^ percentile centred around *r* = 0. **b.** Mean High synchrony (HS) vs. low synchrony (LS) in each physiological measure and concert, with standard error. One musical bar is preceding bars correlation with high ISC value starting from the first bar of correlation (bar0) to last bar of correlation window (bar4).

### Acoustic properties as predictors of audience synchrony

#### Tempo

Logistic regression revealed that RR synchrony was significantly predicted by tempo for all three concerts in Beethoven (C1: β = 0.011, C2: β = 0.016; C3: β = 0.007, all *p* < .001) and in Brahms (C1: β = 0.018; C2: β = 0.028; C3: β = 0.008, all *p* < .001). Figure 3b shows that faster tempo increased probability that RR was highly synchronised across audience members. Probability of SCR synchrony significantly increased by faster tempo in Brahms for all concerts (C1: β = 0.028; C2: β = 0.036; C3: β = 0.015, all *p* < .001) and in Beethoven for two concerts (C1: β = 0.007, *p* < .001; C2: β = 0.011, *p* < .001; C3: β = 0.001, *p* = .61). HR and EMG synchrony was not consistently predicted by tempo. Probability of synchronised HR increased at slower tempi in Beethoven C2 (β = −0.009, *p* < .001), but at faster tempi in Brahms C3 (β = 0.007, *p* =.018). Probability of EMG synchrony decreased with faster tempi in Beethoven C2 (β = −0.006, *p* = .019), but increased in Dean C3 (β = 0.024, *p* = .003).

**Figure 3.**
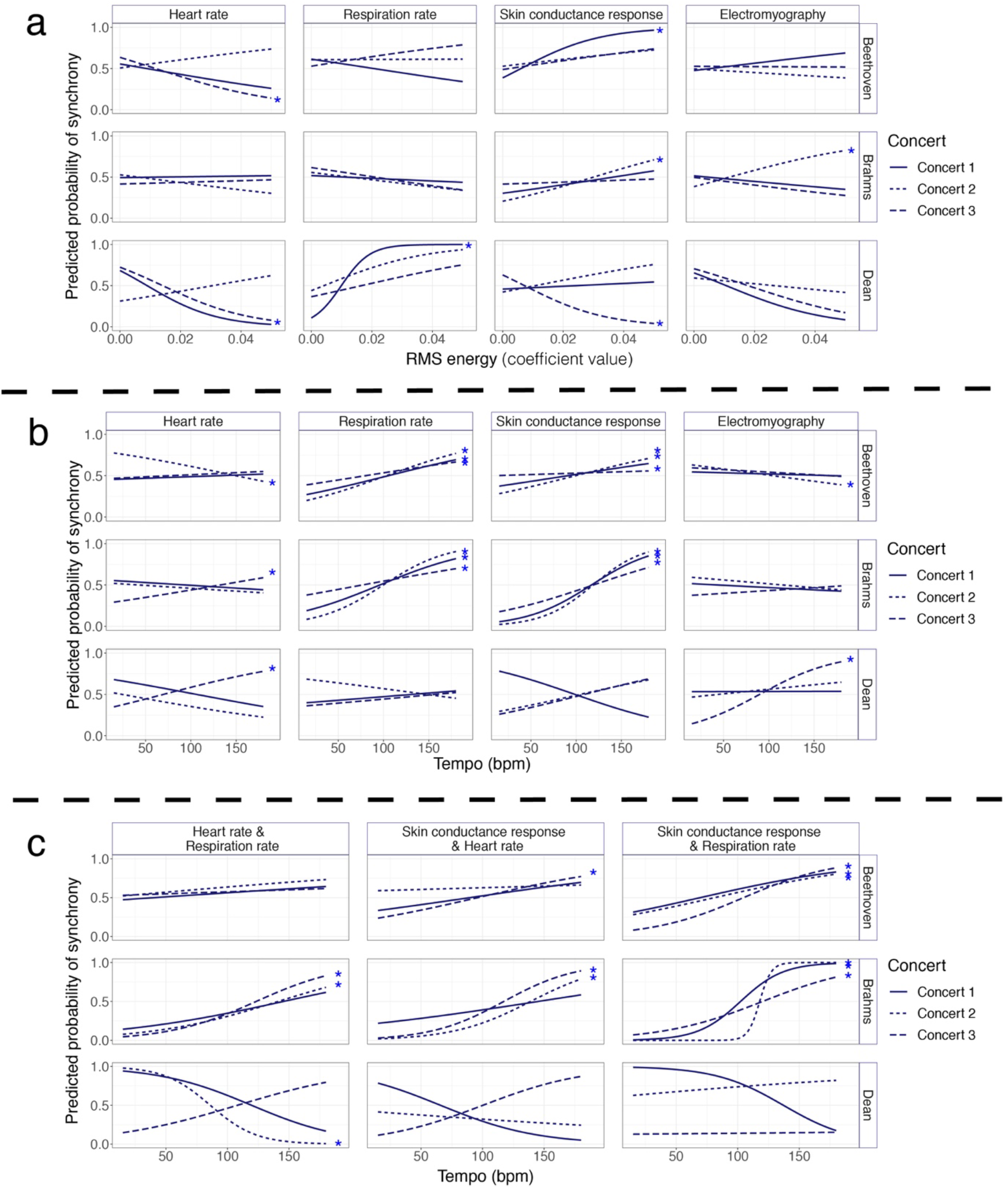
Probability curves of high (1) vs. low (0) synchrony across listeners extracted for individual acoustic predictors in the logistic regression models. Separate panels show individual acoustic predictors (**a**: RMS; **b** and **c**: tempo) for each physiological measure (per column), in each piece (per row in each panel) and concert (indicated by line style).

#### RMS energy

Figure 3a shows that lower RMS energy increased probability of HR synchrony, but this was not consistent across concerts or pieces and only significant in Beethoven C3 (β = −47.39, *p* = .002) and Dean C1 (β = −86.00, *p* = .033). Higher RMS energy significantly increased probability of SCR synchrony in the Classical/Romantic works, but not consistently across concerts, i.e., only significant for Beethoven C1 (β = 76.20, *p* < .001) and Brahms C2 (β = 44.69, *p* = .007). Higher RMS increased probability of RR synchrony in Dean C1 (β = 212.80, *p* < .001) and EMG synchrony in Brahms C2 (β = 40.17, *p* = .006).

#### Spectral centroid

Probability of HR synchrony increased with higher spectral centroid in Beethoven C2 (β = 0.001, *p* = .009), but with lower spectral centroid in Beethoven C3 (β = −0.001, *p* = .029) and in Dean C1 (β = −0.001, *p* = .048). Lower spectral centroid significantly increased probability of RR synchrony in Beethoven C2 (β = −0.001, *p* = .047) and Dean C2 (β = −0.001, *p* = .049). Higher spectral centroid increased probability of SCR in Dean C2 (β = 0.002, *p* = .006) and probability of EMG synchrony in Brahms C2 (β = 0.001, *p* = .026) and C3 (β = 0.001, *p* =.0018) and in Dean C3 (β = 0.002, *p* = .021).

#### Key clarity

Lower key clarity increased probability of HR synchrony in Beethoven C3 (β = −2.09, *p* = .050), of RR synchrony in Dean C1 (β = −5.71*, p* = .007), and of SCR synchrony in Dean C1 (β = −4.67, *p* = .009).

#### Derivatives of MIR features

No consistent predictors were observed. Tempo changes predicted SCR synchrony in Dean C3 (β = −0.02, *p* = .044). Spectral centroid changes predicted HR synchrony in Brahms C1 (β = −0.001, *p* = .032) and SCR synchrony in Beethoven C2 (β = 0.001, *p* = .025).

### Synchrony across multiple physiological measures

As SCR, HR, and RR are all responses of the ANS, we sought to asses which musical features may predict an unified ANS response, i.e., when all three physiological measures were in synchrony simultaneously (when HS bars were the same in SCR, HR, and RR). Unfortunately, synchrony of the ANS as one entity (SCR, HR, RR) was not possible to model as no LS moments were found in the Dean piece for C1. Splitting ANS responses into paired combinations yielded HS and LS moments in all three styles in all three concerts allowed further modelling.

Faster passages (around 120 bpm, see Figure 3c) significantly increased probability of combined SCR-RR synchrony for all concerts in Beethoven (C1: β = 0.014, *p* = .039; C2: β = 0.014, *p* = .031; C3: β = 0.027, *p* = .013) and Brahms (C1: β = 0.058, *p* < .001; C2: β = 0.212, *p* = .002; C3: β = 0.025, *p* = .032), but not in Dean (all *p* > .4). Faster tempi increased probability of combined SCR-HR synchrony in Beethoven C3 (β = 0.014, *p* = .019), and Brahms C2 and C3 (C2: β = 0.032, *p* = .027; C3: β = 0.034, *p* = .004). Faster tempi increased probability of HR-RR synchrony in Brahms C3 (β = 0.028, *p* = .012). Slower tempi increased probability of HR-RR synchrony, but only for Dean C2 (β = −0.054, *p* = .018). RMS only occasionally predicted combined physiological synchrony, where higher RMS increased probability of combined SCR-RR synchrony in Beethoven C1 (β = 82.876, *p* = .038) and marginally in Dean C1 (β = 538.170, *p* = .054). It also marginally increased probability of SCR-HR synchrony in Beethoven C2 (β = 71.00, *p* = .057). Spectral centroid and key clarity rarely predicted combined physiological synchrony. Higher spectral centroid increased the probability of SCR-RR synchrony in Beethoven C3 (β = 0.002, *p* = .001) and SCR-HR synchrony in Brahms C3 (β = 0.004, *p* = .014). Lower key clarity increased probability of HR-RR synchrony in Brahms C1 (β = −11.48, *p* = .042) and SCR-RR synchrony in Dean C1 (β = −19.19, *p* = .044).

### Physiology synchronisation across concerts to higher level features

As we used naturalistic music, it was important to consider stylistic and compositional features of the music which are not easily analysed computationally. Therefore, we used standard music theoretical approaches^77,78^ to investigate higher-level musical events of the most ‘salient’ moments, operationalised by two criteria: when 1) high physiological synchrony in any of the physiological measures occurred in all three concert audiences and 2) sustained synchrony was for more than one bar.

Overall, audience physiology seemed to synchronise around three types of musical events: a) transitional passages with developing character; b) clear boundaries between formal sections; and c) phrase repetitions (all listed with descriptions in Supplementary Table S3). Salient responses occurred during ‘calming down’ (e.g., Beethoven 1^st^ movement, [Beethoven1], bars [b] 85-88; b287-293; Dean2, b70-71; Dean4, b75-77; Brahms4, b75-76) or arousing (e.g., Beethoven1, b303-307; Dean2, b23-25; Brahms3, b6-8) transitional passages, characterised by a decrease or an increase of loudness, texture, and pitch register respectively. Other salient responses occurred when there was a clear boundary between functional sections, indicated through parameters such as a key change (e.g., between major and minor key in Beethoven3, b84-88; Brahms3, b58-61), a tempo change (e.g., Beethoven 1 b328-331; Brahms4, b248-250), or a short silence (e.g., Beethoven1, b96-97; Beethoven4, b10-14). Lastly, salient responses occurred when a short phrase or motive was immediately repeated in a varied form, for example in an unexpected key, (e.g., Beethoven1, b35-37), elongated or truncated (e.g., Beethoven1, b85-88; 291-293), or with a different texture or pitch register (Brahms1, b90-91; Brahms3, b170-171). Since the immediate varied repetition of a short phrase is very common in Classical and Romantic styles, salient responses were also evoked when a phrase repetition occurred simultaneously with a transition or clear boundary (e.g., Beethoven1, b24-30; b136-138). With regard to style, the three categories are in line with the compositional conventions of the respective works: salient responses were found more often during transitions in the Romantic and Contemporary works, and during phrase repetitions and boundaries in the Classical work.

## Discussion

We demonstrate a novel approach to implicitly assess the continuous group music listening experience in a naturalistic environment using physiological ISC and inter-disciplinary stimulus analysis. By measuring physiological responses of audiences listening to live instrumental music in a typical concert setting, we examined which musical features evoked synchronised responses (operationalised via ISC). Consistency of effects was assessed by repeating the same concert three times with different audiences. Importantly, we found no significant differences of length, loudness, tempo, or timbre across the concert performances, allowing us to assume that varying patterns of audience responses were due to the weakness of an effect or individual differences in audiences rather than a difference in acoustics. When responses were highly correlated (within a 5-bar window), there was an overall higher SCR and EMG (smiling muscle) activity, and increasing HR and RR, suggesting that synchronised physiology was associated with increased arousal.

Synchronised RR and SCR responses were consistently predicted by tempo alone. Additionally, tempo predicted not only individual physiological measures, but a more general ANS response, that is, when both SCR and RR of audience members became synchronised simultaneously. This finding suggests further that tempo induces reliable responses, in line with previous work showing that tempo and rhythm are the most important musical features in determining physiological responses^31^.

The current results show that faster tempi consistently increased probability of combined SCR-RR synchrony. As faster tempo is typically perceived as more arousing^31–34^, our finding supports previous research linking high ISC to higher arousal^59,61,72^. These results could further support the idea that ISC is related to stimulus engagement^5^: as slower music increases mind-wandering^79^, slower tempi may result in reduced attention to the music, leading to greater individual variability in physiological responses and subsequently lower ISC^59,67^. However, we wish to note that faster tempi (centred around 120 bpm or 2 Hz in the current study) might also be a more physiologically optimal and/or perceptually familiar range for entrainment to music (see^80^ for review), compared to slower tempi (centred around 50 bpm .83 Hz in the current study). Entrainment, or perhaps just adaptation of a physiological measure towards the musical tempo, might be one mechanism through which faster (or optimally resonant) tempi induce more similar audience responses.

It is of further interest that SCR-RR synchrony was more probable at faster tempi only in the Classical and Romantic styles, but not in the Contemporary style. We suggest, therefore, that the effect of tempo may be modulated by the context in which it occurs, supporting previous studies showing that the same features evoke different physiological responses based on the style^38,51^ and/or the familiarity and engagement with the music^5^. It is worth noting that while Beethoven and Brahms were rated as more familiar compared to Dean, reported engagement did not differed across styles (see^81^). This suggests that familiarity of style, rather than engagement, may increase the probability of synchrony (though this difference could be due to our stimuli being presented live rather than through a recording^5^). Beethoven and Brahms have a relatively structured and stable meter with very few instantaneous tempo changes within movements, whereas many Dean passages contain unstable meter (e.g., the first movement has alternating bars of four, five, or six beats) and frequent tempo changes within movements (as it typically for each style). In view of this, the fact that synchrony probability changes between styles could also be due to stimuli coherence. This reasoning is in line with the idea of higher ISC occurring in more predictable contexts^67^ and lower ISC occurring in versions of music where the beat was disrupted^70^. However, such interpretation may be limited by the fact that we presented only one work per style and tempo was not evenly represented across styles, though this compromise was dictated by the constraints of a naturalistic concert setting. Nonetheless, this suggests that tempo may be a driving aspect in predicting synchronised physiological responses; though further research would be required to assess whether coherence^70^ and/or engagement/familiarity^5^ are modulatory mechanisms of synchrony.

Although orienting/startle response research consistently shows that loudness evokes highly replicable physiological responses in a controlled tone sequence^23,26,27^, our results suggest that synchronised physiological responses across concert audiences are not consistently statistically predicted by RMS energy, nor by key clarity and spectral centroid (and neither changes of these features) in the music. This finding may point to the importance of context: previous work has shown that environmental sounds and music are experienced physiologically differently, such as an increase in HR (index of a startle response^21^) with arousing noises (e.g., a ringing telephone or storm), but not with music^82^. As loudness in the current study was embedded in naturalistic music (rather than in a tone sequence), this highlights the generalisability limitations of reductionist stimuli to real-world contexts^54^. However, this points to another limitation: despite employing such complex stimuli, we computationally extracted only the most common acoustic parameters associated with physiological responses, possibly leading to underfitting of our models^83^. It was important, therefore, to explore additional higher-level parameters in the music that are not so easily extracted computationally.

We observed that transitional passages, clear boundaries or immediate phrase repetitions in the music – identified using music theoretical analysis – coincided with highly synchronised physiological responses across concerts. This observation corroborates previous findings that synchrony may occur in response to long-term, structural features in music^4,70^. The fact that synchronised responses were evoked by arousing transitional passages (characterised by changes in loudness, pitch register, and musical texture) supports the idea that highly similar responses in all audiences are related to arousal^59,61,72^ as well as supporting the idea that audience members collectively ‘grip on’ to loudness and texture changes^6^. As unexpected musical events embedded in a predictable context may be perceived as arousing^12,46,47^, our findings that physiological synchrony occurred during sudden tempo or key changes (i.e., clear musical event boundaries), further support the notion that correlated responses occur at arousing moments^59,61,72^. This finding aligns with the idea that disruptions of temporal expectations affect ANS responses^27,84^ as well as synchrony in EEG components^70^. Additionally, or alternatively, it is likely that surprising events phase-reset ongoing physiological oscillations (see^85,86^ for reviews), perhaps thereby leading, at least briefly, to an increase in audience synchrony around moments of phase resetting. The finding that synchronised responses occurred during immediate phrase repetitions hint a general attention towards repetitions in music^87^. Regarding the recurrence of phrases after longer intervals, it remains unclear if audience synchrony reflects an ‘orientation response’^20,21,27^, indicating that they recognize thematic connections over larger time spans^88^. Rather, our analysis suggests, that an interplay of various musical features, in addition to the simple repetition, increase attention of all audience members to these musical moments and subsequently enhance audience synchrony. For instance, high collective audience synchrony occurred in some of the structurally most important moments of Beethoven1, such as the end of the exposition (b136-138: phrase repetition and boundary), deferred cadences (declined structural closures in b96-97, b301-302: boundary), and references to the main theme at the end of the movement (b328-331: boundary). Although ISC has been found to decrease with repeated listening of the same piece (if in a familiar style)^5^, the fact that high ISC occurred at unexpected phrase repetitions is in line with compositional practices, in which a composer tries to vary and develop thematic material^89^ (with a different texture or harmonically) to keep ‘interest’. Although this part of our analysis remains observational and exploratory, it nonetheless points to certain musical features which could be systematically manipulated in future studies.

In conclusion, by measuring continuous music listening experience in a naturalistic setting of a chamber music concert, we show that synchronised physiological responses across audience members (operationalised via ISC) are predicted by tempo and may be linked to structural transitions, boundaries, and phrase repetitions. Our results support the idea that group synchronisation is linked to musical arousal, structural coherence, and familiarity. Using naturalistic music in such a concert environment is beneficial in that participants are more likely to be absorbed in the music^6^ and have more realistic and stronger responses^8,9^. However, this makes our findings specific to the music we have used, especially as we utilised only one piece per style. Future research should assess whether the current findings related to musical features and style are replicated with different kinds of music, both within and outside of the styles used in this study, as well as a wider range of musical features to improve characterisation of such complex stimuli. Further questions remain for the concert setting itself; for example, whether these effects and perceptions would change with and without visual information of the performer, or with varying programming orders, performance spaces, and concert aspects^90^. Exploring musical experiences from pre-recorded or live performances – with or without the co-presence of others – may prove an interesting future research direction, especially with regard to the COVID-19 pandemic and current transformations of the live concert experience.

## Method

**Participants, materials, and experimental procedure** are identical to Merrill et al.^81^. All experimental procedures were approved by the Ethics Council of the Max Planck Society, and were undertaken with written informed consent of each participant. 138 participants attended one of three evening concerts (starting at 19.30 and ending at approximately 21.45) in a hybrid performance hall purpose-built for empirical investigations (the ‘ArtLab’ in Frankfurt am Main, Germany). Care was taken to keep parameters (e.g., timing, lighting, temperature) as similar as possible across concerts. Professional musicians performed string quintets in the following order: Ludwig van Beethoven, op. 104 in C minor (1817), Brett Dean, ‘Epitaphs’ (2010), a 20-minute interval, Johannes Brahms, op. 111 in G major (1890). Continuous blood volume pulse (BVP), respiration data, skin conductance, and facial electromyography (EMG) from the zygomaticus major muscle were measured with a plux device (https://plux.info/12-biosignalsplux) for the entirety of the concert at 1000 Hz. After excluding participants with more than 10% missing data^91^, physiological data from 98 participants (that had comparable education levels and age distribution across concerts) and acoustic data from musical recordings were pre-processed and analysed in MATLAB 2018b.

### Data analysis

#### Musical feature extraction

Using Sonic Visualiser^92^, instantaneous tempo was manually extracted by tapping each beat, calculating inter-onset intervals (IOIs) between each beat, and then converting to beats per minute (bpm). All other features were computationally extracted using the MIRToolbox^93^. RMS energy (related to loudness), spectral centroid, brightness, and roughness (related to timbre) were extracted using 25 ms windows with 50% overlap^94^. Key clarity was extracted using a 3 second window with 33% hop factor^1^. As previous time-series analyses have parsed data into meaningful units of clause and sentence lengths^64^, a meaningful unit in music is a bar (American: measure). Correspondingly, values were aligned by averaging each feature into bins per bar (on average 10 seconds). It is worth noting that acoustic features can be distinguished between compositional features and performance features^95^, where the former are represented in the musical score (such as harmony), and the latter include features that can change between performances, namely how loud and fast musicians may perform the music. Because key clarity is a compositional feature (i.e., does not change between performances), we averaged values across concert performances.

When checking for independence of features^96^, RMS and roughness correlated highly (*r* > .7) as did brightness and spectral centroid (*r* > .7) in all movements. As RMS and spectral centroid are features more commonly used compared to roughness and brightness^95,97^, and spectral centroid seems to best represent brightness^75^ and overall timbral^98–100^ perception, we kept only key clarity, RMS, spectral centroid, and tempo. The degree of change in these features was also obtained, that is, the difference between adjacent bars. To compare acoustic features per style, linear mixed models with fixed effect of the works (Beethoven, Brahms, and Dean) and random effect of movement were constructed per acoustic feature and per concert. To check performance feature similarity between concerts, each feature was compared with concert (C1, C2, C3) as the independent variable. Pearson correlations were used to assess similarity of acoustic features over time between concerts (C1-C2, C1-C3, C2-C3). Correlations (false-discovery rate corrected) were considered adequate if they met a large effect size of concert *r* > .5^101^.

#### Physiology pre-processing

Data were cut per movement. Missing data (gaps of less than 50ms) were interpolated at the original sampling rate. Fieldtrip^102^ was used to pre-process BVP, respiration, and EMG data. BVP data were band-pass filtered between 0.8 and 20 Hz (4^th^ order, Butterworth) and demeaned per movement. Adjacent systolic peaks were detected to obtain inter-beat intervals (IBIs) and an additional filter was added to remove any IBIs that were shorter than 300 ms, longer than 2 seconds, or had a change of more than 20% between adjacent IBIs (typical features of incorrectly identified IBIs^103^). After visual inspection and artefact removal, IBIs were converted to continuous heart rate (HR) by interpolation. Respiration data were low-pass filtered (.6Hz, 6th order, Butterworth) and demeaned. As in the BVP data, maximum peaks were located and respiration rate (RR) was inferred by the peak intervals. EMG activity was bandpass filtered (between 90 and 130 Hz, 4^th^ order, Butterworth), demeaned and Hilbert transformed. Skin conductance data were pre-processed using Ledalab^14^ and decomposed into phasic and tonic activity. As we were interested in event-related responses, only (phasic) skin conductance responses (SCR) were used in further analyses. All pre-processed physiological data (SCR, HR, RR, EMG) were resampled at 20Hz^19^, *z*-scored within participant and movement, and averaged into bins per bar.

#### ISC analysis

We calculated a time-series ISC based on Simony et al.^67^ by forming *p* × *n* matrices (one for each SC, HR, RR, and EMG, and for each of the twelve movements per concert), where *p* is the physiological response for each participant over *n* time points (bars across the movements) over a sliding window 5 bars long (approximately 10 seconds; the average bar length across the whole concert was 2 seconds), shifting one bar at a time. Fisher’s *r*-to-*z* transformation was applied to correlation coefficients per subject, then averaged *z* values were inverse transformed back to *r* values. The first 5 bars and the last 5 bars of each movement were discarded to remove common physiological responses evoked by the onset/offset of music^65^. ISC values per movement were concatenated within concerts (2238 bars, see Figure 2a), giving four physiological ISC measures per concert. These ISC traces represent the similarity of the audience members’ physiological responses over time.

Following Dmochowski et al.^61^, bars of high/low synchrony (HS/LS) were defined using 20^th^ percentiles. Physiological response that were highly synchronised (HS) across audience members were defined as bars where ISC values rose above an 80-percentile threshold, while low synchrony (LS) was defined as ISC values within a 20-percentile centred around zero, that is, with low correlation. To obtain instances of overall ANS synchrony (i.e., across multiple physiological measures simultaneously), we identified where HS and LS moments of one physiological measure coincided at the same time as another physiological measure. Physiological responses at points of HS and LS were compared using linear models with factors Synchrony (HS, LS) and Bar (bars 0-4). To investigate whether acoustic features predicted HS vs. LS of physiological responses across audience members, tempo, RMS energy, key clarity, and spectral centroid in bars of HS and LS were recovered. By dummy-coding Synchrony as a binary variable, with HS as 1 and LS as 0, logistic regression models were constructed to predict Synchrony for each physiological measure (dependent variable) with continuous predictors of tempo, key clarity, loudness, and spectral centroid from the HS and LS bars (all features were included, as perceived expression in music tends to be determined by multiple musical features^34^). (N.B.: no random intercept of movement was included, because not all movements contained epochs of significantly high or low ISC). As we expected style to modulate the effect of these acoustic features in predicting synchrony, models were run separately per piece and concert.

#### Statistical analyses

were conducted in R^104^. Pearson correlations were computed using *corr.test* in the *psych* package and adjusted for false discovery rate using the Benjamin-Hochberg procedure. Linear models and linear mixed effects models were constructed using the *lme4* package^105^; *p* values were calculated with the *lmerTest* package^106^ via the Satterthwaite approximation and using the Anova function in the *car* package^107^. Contrasts were assessed with the emmeans function (*emmeans* package^108^), with the Tukey method of adjustment. Logistic regressions models where run using a general linear model with a logit link function.

#### Music theoretical analysis

The scores of all works were analysed according to widely used methods for the respective styles (Classical and Romantic: Hepokoski & Darcy^77^; Contemporary: Zbikowski^78^ and Schoenberg^89^). Musical events were labelled on the beat level (harmonic changes, cadences, texture changes, etc.) and larger sections (e.g., transition between primary and secondary action space) according to the convention of the previous methods. The performance recordings and acoustic results served as reference for passages which could have been interpreted equivocally in the score. After the analysis, passages involving high physiological synchrony were marked and categorized according to common features across styles; these features where then deduced into appropriate categories.

## Acknowledgments

Many thanks go to staff at the ArtLab (Frankfurt am Main), in particular Alexander Lindau, Patrick Ulrich, and Eike Walkenhorst who helped organise the concerts; to Claudia Lehr and Freya Materne for organizing the invitations of the participants, and finally, thanks to the many assistants during data collection, particularly Till Gerneth, Sandro Wiesmann, Nancy Schön, and Simone Franz. We would like to thank Folkert Uhde, as part of the Experimental Concert Research (ERC) project, who set up the musical program for this concert. Thanks also to Cornelius Abel and Alessandro Tavano for data quality checks; to Helen Singer for annotating musical beats, and Elke Lange for advice on statistics and MIR feature extraction.

## Author contributions

**A.C.** design of the work; statistical and musicological analysis; data interpretation; writing – original draft and figures. **L.K.F.** design of the work; statistical analysis; data interpretation; writing – review and editing. **L.T.F.** musicological analysis; writing – review and editing. **M.T.** conception. **M.W.F.** conception; data acquisition; data interpretation; writing – review. **J.M.** design of the work; data acquisition and statistical analysis; data interpretation; writing – review and editing.

## Additional Information

### Data availability

Data of this study are available from the corresponding author upon request.

