## Supplementary Table S1, 2 and 3 for "Synchrony in the periphery: inter-subject correlation of physiological responses during live music concerts"

### Supplementary Information

*Supplementary Table S1a. F–tests for performance (acoustic) differences across concerts.*

| | | Beethoven | | Brahms | | Dean | |
|---|---|---|---|---|---|---|---|
| | df | F | *p* | F | p | F | *p* |
| RMS | 2 | 2.14 | 0.12 | 0.23 | 0.79 | 0.12 | 0.89 |
| Spectral centroid | 2 | 1.78 | 0.17 | 1.54 | 0.21 | 2.82 | 0.06 |
| Tempo | 2 | 2.56 | 0.08 | 0.02 | 0.98 | 2.31 | 0.10 |

*Supplementary Table S1b. Lengths of separate movements of works.*

| | Length in minutes per movement (mvt) | | | | | | | | | | | |
|---|---|---|---|---|---|---|---|---|---|---|---|---|
| | Beethoven | | | | Dean | | | | Brahms | | | |
| | 1st mvt | 2nd mvt | 3rd mvt | 4th mvt | 1st mvt | 2nd mvt | 3rd mvt | 4th mvt | 1st mvt | 2nd mvt | 3rd mvt | 4th mvt |
| C1 | 7.80 | 8.15 | 3.76 | 6.37 | 2.53 | 3.87 | 3.90 | 3.39 | 10.17 | 7.15 | 5.37 | 4.94 |
| C2 | 7.95 | 8.36 | 3.74 | 6.50 | 2.70 | 3.87 | 3.93 | 3.49 | 10.24 | 7.07 | 5.39 | 4.95 |
| C3 | 7.75 | 8.02 | 3.71 | 6.23 | 2.68 | 3.87 | 3.79 | 3.14 | 10.03 | 6.97 | 5.46 | 4.93 |

*Supplementary Table S2. F–tests for linear models comparing physiology in 5 bar windows for Synchrony (HS / LS) across correlation window in terms of Bar (0-4).*

| | | HR | | | | | | RR | | | | | |
|---|---|---|---|---|---|---|---|---|---|---|---|---|---|
| | | C1 | | C2 | | C3 | | C1 | | C2 | | C3 | |
| | df | F | *p* | F | *p* | F | *p* | F | *p* | F | *p* | F | *p* |
| Synchrony | 1 | **13.31** | **< .001** | **24.43** | **< .001** | **10.07** | **0.002** | **5.15** | **.023** | **8.05** | **.005** | **25.53** | **> .001** |
| Bar | 4 | **4.14** | **0.002** | 1.74 | .13 | **3.74** | **0.004** | **5.49** | **< .001** | **10.26** | **< .001** | **10.70** | **> .001** |
| Synchrony x Bar | 4 | **3.34** | **.010** | 0.85 | 0.50 | **3.04** | **0.016** | 2.19 | 0.068 | **6.30** | **< .001** | **10.35** | **> .001** |
| | | SCR | | | | | | EMG | | | | | |
| | | C1 | | C2 | | C3 | | C1 | | C2 | | C3 | |
| | df | F | *p* | F | *p* | F | *p* | F | *p* | F | *p* | F | *p* |
| Synchrony | 1 | **103.22** | **< .001** | **52.67** | **< .001** | **90.17** | **< .001** | **20.04** | **< .001** | **12.82** | **< .001** | **29.24** | **< .001** |
| Bar | 4 | **28.97** | **< .001** | **26.98** | **< .001** | **24.78** | **< .001** | **5.34** | **< .001** | **8.04** | **< .001** | **6.41** | **< .001** |
| Synchrony x Bar | 4 | **26.39** | **< .001** | **19.37** | **< .001** | **15.98** | **< .001** | **4.76** | **< .001** | **5.97** | **< .001** | **7.85** | **< .001** |

*Supplementary Table S3a. Musical descriptions of bars with salient physiological responses: Beethoven*

| Ludwig van Beethoven: String Quintet in C minor op. 104 (1817) | | | |
|---|---|---|---|
| **Movement** | **Bars** | **Categories** | **Description** |
| 1st * | 24-30 | Phrase repetition<br>Transition | Exposition: Increasing texture, dynamics & harmonic rhythm, modulation and half-cadence bar30 ending the primary-theme zone (P) |
| | 35-37 | Phrase repetition | Exposition: Chromatic sequence at opening of transition (T) |
| | 85-88 | Phrase repetition<br>Transition | Exposition: Decreasing texture & dynamics, bar-wise repetition of motif, elongation & prolongation of secondary diminished dominant |
| | 96-97 | Boundary | Exposition: Strong half-cadence, essential expositional closure (EEC) declined in b97 |
| | 136-138 | Phrase repetition<br>Boundary | Exposition: Increasing texture & dynamics, syncopations, cadence ending the closing zone (C) of exposition. First chord of development section. |
| | 287-290 | Phrase repetition<br>Transition | Recapitulation: Decreasing texture & dynamics, bar-wise repetition of motif, prolongation of secondary diminished dominant |
| | 291-293 | Phrase repetition | Recapitulation: Elongation & prolongation of secondary diminished dominant |
| | 301-302 | Boundary | Recapitulation: essential structural closure (ESC) declined in b300, beginning of strong transitional section, motivic development of P in unison |
| | 303-307 | Transition | Recapitulation: strong transitional section, motivic development of P in unison |
| | 328-331 | Boundary | Coda: Reference to opening bars with fermata & tempo change (150bpm to Adagio, ca. 80 bpm and back) after final cadence in b327 |
| 2nd | 62-64 | Boundary | Variation 1: Perfect authentic cadence (PAC), end of var 1 |
| | 126-130 | Transition<br>Boundary | Variation 3: increase of dynamic (*morendo*), PAC b129 ending var. 3 in minor.<br>Variation 4: Key, texture, tempo change, beginning of var.4 in major |
| 3rd | 65-67** | Phrase repetition | Menuetto 1, B-part 2nd time: standing on the dominant after half cadence (HC) b64, reference to beginning of B-part b55-58 |
| | 84-88 | Boundary | Menuetto 1, A'-part 2nd time: end of menuetto (minor), beginning of Trio (major). Change of key, register, texture, phrase rhythm |
| 4th * | 10-14 | Boundary<br>Transition | Exposition: beginning of primary theme one (P1) after intro & general rest in b8 |
| | 150-153 | Phrase repetition<br>Boundary | Development: continuation within P0 at the beginning of development,<br>half cadence, general rest |
| | 187-188 | Phrase repetition<br>Boundary | Development: rotation of subordinate theme (S), second pair of of six-bar chromatic sequence of S |

*Note: no repeat. **Note: Bars in relation to repeats

*Supplementary Table S3b. Musical descriptions of bars with salient physiological responses: Dean*

| Brett Dean: Epitaphs (2010) | | | |
|---|---|---|---|
| **Movement** | **Bars of high synchrony** | **Categories** | **Description** |
| 2nd | 23-25 | Transition | Increasing texture density, rhythmically clearer than previous context |
| | 61-62 | Transition<br>Boundary | Closing phrase with decreasing loudness<br>cadence and *finalis* with fermata |
| | 70-71 | Transition | Increasing loudness and dissonance, varied repetition, change of pitch class quality & timbre (*increasingly raw*) |
| 4th | 75-77 | Boundary<br>Transition | Sudden decrease of dynamics, change of texture & timbre<br>Glissando over 3+3+2 pattern |

*Supplementary Table S3c. Musical descriptions of bars with salient physiological responses: Brahms*

| Johannes Brahms: String Quintet in G major op. 111 (1890) | | | |
|---|---|---|---|
| **Movement** | **Bars of high synchrony** | **Categories** | **Description** |
| 1st * | 90-91 | Phrase repetition | Development: combined P/S space, homophonic texture |
| 3rd | 6-8 | Transition | 1st minor part, A: modulation to half cadence in the middle of 8-bar continuation. Small increase of dynamics, descending line in upper melody |
| | 58-61** | Boundary | 1st minor part, B: end of first, beginning of second repeat. Final cadence with *finalis* b58 lowest pitch of the melody, change to major b59, change of register and texture b61 |
| | 170-171** | Boundary<br>Phrase repetition | 2nd minor part, A: end of 12-bar antecedent on HC, beginning of consequent (repeat of antecedent), change of register to upper octave |
| | 218-219 | Boundary | 2nd major part, coda: end of 2nd minor part, beginning of coda, change of key |
| 4th | 75-76 | Transition | Exposition: within coda, change of texture & timbre, decrease of dynamics, modal cadence with 5-6 progression |
| | 248-250 | Boundary | Coda: beginning of *stretta* (120 bpm to 140 bpm), homophonic, clear rhythm |

*Note: no repeat. **Note: Bars in relation to repeats
